# Secretory Autophagy via VMP1-Containing Extracellular Vesicles in Pancreatic Stress Responses

**DOI:** 10.1101/2024.10.31.615473

**Authors:** Mariana S. Tadic, Felipe J. Renna, Malena Herrera López, Fabiana López Mingorance, Tamara Orquera, Diego A. Chiappetta, Alejandro Ropolo, Maria I. Vaccaro

## Abstract

Cellular stress activates mechanisms such as autophagy and vesicular trafficking to maintain homeostasis in pathological conditions like acute pancreatitis. Vacuole Membrane Protein 1 (VMP1), an autophagy-related protein implicated in pancreatitis, diabetes, and pancreatic cancer, triggers autophagy through ubiquitination and interaction with BECN1. Here, we show that VMP1 is secreted into the extracellular medium and incorporated into extracellular vesicle (EV) membranes. Using cells expressing VMP1-tagged plasmids, we isolated VMP1- containing EVs (VMP1-EVs) by ultracentrifugation and immunoisolation. VMP1-EV secretion decreased with mTOR inhibition and in Atg5-deficient cells. In pancreatic acinar cells, endogenous VMP1 secretion increased under stress, including blocked autophagic flux and experimental pancreatitis. In a rat pancreatitis model, VMP1 secretion in pancreatic juice was also elevated. TEM and DLS analyses revealed VMP1-EVs of ∼150 nm. LC3-II was detected in VMP1-EVs, and its release increased under lysosomal blockade. VMP1 downregulation reduced LC3 and p62 secretion, demonstrating that VMP1 drives a secretory autophagy pathway relevant to pancreatic pathophysiology.

## 1. Introduction

Macroautophagy/autophagy is a tightly regulated catabolic process that mediates the selective degradation of damaged or superfluous cellular components, thereby preserving cellular homeostasis and promoting survival under stress [1]. While autophagy is classically regarded as a degradative pathway, accumulating evidence has expanded its functional repertoire to include non-canonical roles in secretion. This has led to the recognition of secretory autophagy (SA) [2], a process in which autophagic components are repurposed for unconventional secretion, encompassing the release of leaderless cytosolic proteins, secretory lysosomes, and extracellular vesicles (EVs) [2,3]. EVs are membrane bound vesicles secreted by cells into the extracellular space. EVs exhibit diverse sizes, contents, and surface markers [4,5].

Despite advances in our understanding of SA, the precise molecular mechanisms underlying this process remain incompletely defined. Autophagy and endosomal trafficking are mechanistically intertwined, sharing membrane sources, regulatory proteins, and vesicular dynamics. Autophagy related proteins - including ATG5, LC3 (microtubule-associated protein 1 light chain 3) and SQSTM1/p62 (sequestosome 1). - participate in both degradative and secretory events [6,7]. These intersections highlight the broader role of autophagy beyond lysosomal degradation, extending to vesicle-mediated intercellular communication.

SA is increasingly recognized as a key contributor to diverse pathophysiological conditions, including inflammation, cancer, neurodegeneration, and metabolic diseases [8]. It is typically induced under stress conditions such as nutrient deprivation, endoplasmic reticulum (ER) stress, and accumulation of misfolded proteins that trigger the unfolded protein response (UPR) [9]. Under these circumstances, components normally targeted for lysosomal degradation may instead be secreted through autophagy-dependent mechanisms.

Vacuole Membrane Protein 1 (VMP1) is a 406–amino acid transmembrane protein first identified through the search for genes differentially expressed in pancreatitis [10]. In a series of previous studies, we demonstrated that VMP1 expression alone is sufficient to induce autophagy in mammalian cells under nutrient-rich conditions, through direct binding of its carboxy-terminal peptide to the BH3 domain of BECN1 [11,12]. Briefly, upon expression, VMP1 is ubiquitinated and interacts with BECN1, thereby facilitating the recruitment of the PI3KC3 (Class III phosphatidylinositol 3-kinase) complex I to sites of autophagosome formation at endoplasmic reticulum–plasma membrane contact points. VMP1 subsequently becomes an integral component of the autophagosome membrane [11–14]. Beyond its role in bulk autophagy, VMP1 is critically important in cellular responses to disease. Its expression is undetectable in the healthy human and murine pancreas but is markedly induced in response to pancreatitis [11,15], pancreatic cancer [16–18], and diabetes [19]. Under stress conditions associated with acute pancreatitis (AP), VMP1 expression triggers zymophagy-the selective autophagic degradation of activated zymogen granules [20], as well as mitophagy-the selective removal of damaged mitochondria [21], playing a protective role and avoiding the progression of pancreatitis to severe disease [22]. The regulation of VMP1 is complex, involving transcriptional and post-translational control. We previously described transcriptional programs involving AKT1–GLI3 and E2F1–EP300 that regulate VMP1 expression [16,18], and more recently, identified VMP1 as a target of CRL4/Cdt2-mediated ubiquitination in pancreatic cancer cells, a modification required for its recruitment to the autophagosome [14].

Given VMP1 localization to autophagosomal membranes and its induction under stress, we hypothesized that VMP1 may also participate in SA. In the present study, we provide the first evidence that VMP1 is secreted via EVs and incorporated into their membrane structure. This secretion is regulated by autophagy and is enhanced under stress conditions such as lysosomal inhibition and experimental pancreatitis. The discovery of this alternative secretory pathway may advance our understanding of VMP1 function in health and disease and open new avenues for investigating autophagy-dependent secretion in the context of pancreatic injury and beyond.

## 2. Results

### 2.1. VMP1 is secreted into the cell culture medium via EVs

To determine whether VMP1 is secreted via EVs, CM from HEK293T cells was subjected to differential ultracentrifugation to isolate the EV-enriched pellet. CD63 was used as an EV-fraction marker (Figure 1A). Cells transfected with VMP1.GFP exhibited a clear GFP signal in the EV fraction by immunoblotting, whereas no signal was detected in EVs from cells transfected with an empty GFP vector (Figure 1B).

**Figure 1.**
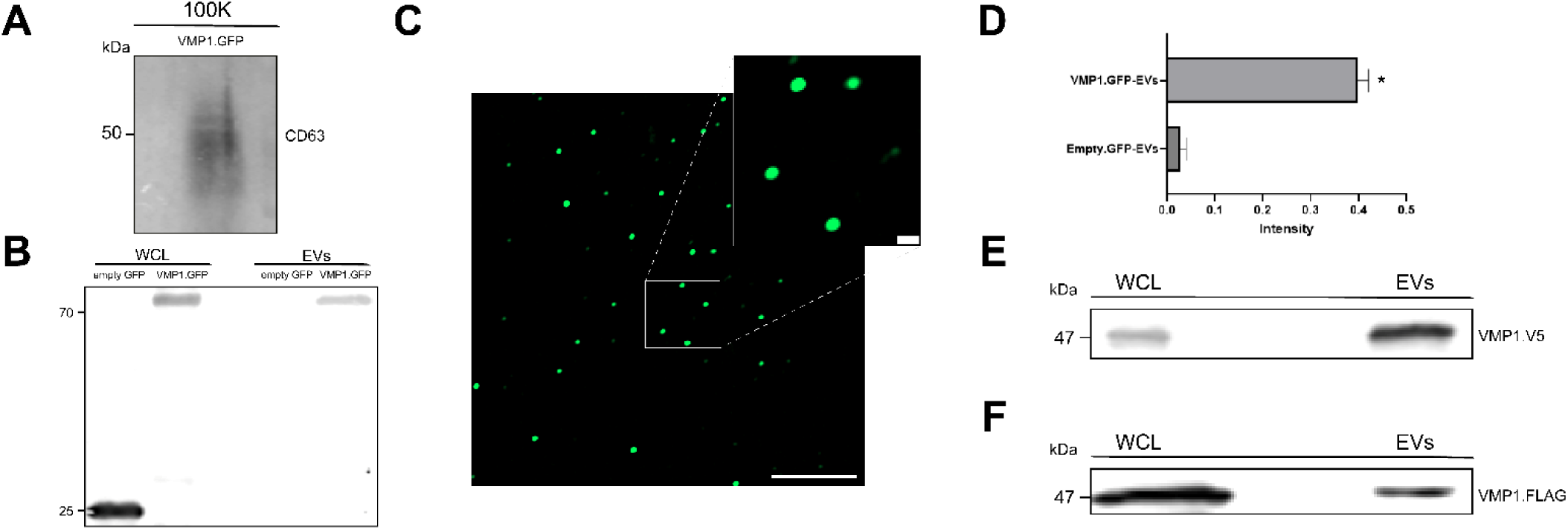
VMP1 is secreted into the cell culture medium via EVs. (A) Detection of CD63 in the VMP1.GFP-EV fraction by immunoblotting. (B) Immunoblot analysis of WCLs and EV fractions isolated by ultracentrifugation from HEK293T cells transfected with VMP1.GFP or empty GFP control. Membranes were probed with an anti-GFP antibody. (C) Representative confocal microscopy images of VMP1.GFP–positive EVs purified from conditioned medium (CM) of HEK293T cells transfected with VMP1.GFP for 48 h. Scale bars: 15 μm (overview), 1 μm (zoomed image). (D) Quantification of GFP fluorescence intensity (FI) in the VMP1.GFP-EV fraction compared to the empty GFP-EV fraction. Data are representative of n = 5 biologically independent experiments for panels A–D, and n = 3 for panel F. (E) Immunoblot analysis of VMP1.FLAG in WCLs and EVs from HEK293T cells using the antibody against FLAG. (F) Detection of VMP1.V5 in WCLs and EVs from HEK293T cells by immunoblotting using anti-V5.

Confocal microscopy of the EV pellet confirmed the presence of VMP1.GFP-positive vesicles with diameters <1 μm (Figure 1C), and fluorescence quantification corroborated enrichment in VMP1.GFP-expressing cells but not in controls (Figure 1D). Similar results were obtained when cells were transfected with VMP1.V5 or VMP1.FLAG constructs, confirming that secretion was independent of the fusion tag (Figures 1E, F). These results demonstrate that VMP1 is secreted into the extracellular space via EVs.

### 2.2. VMP1 is localized to the membrane of VMP1-containing EVs (VMP1-EVs)

Given that VMP1 is a transmembrane protein localized to the autophagosomal membrane, we hypothesized that it retains as a membrane associated protein in EVs. A protease protection assay on EV fractions from VMP1.GFP-expressing cells showed that trypsin treatment of intact EVs abolished the full-length VMP1.GFP signal (∼70 kDa), leaving a ∼25 kDa GFP fragment detectable with anti-GFP antibodies (Figure 2A), indicating C-terminal exposure of GFP on the EV surface.

**Figure 2.**
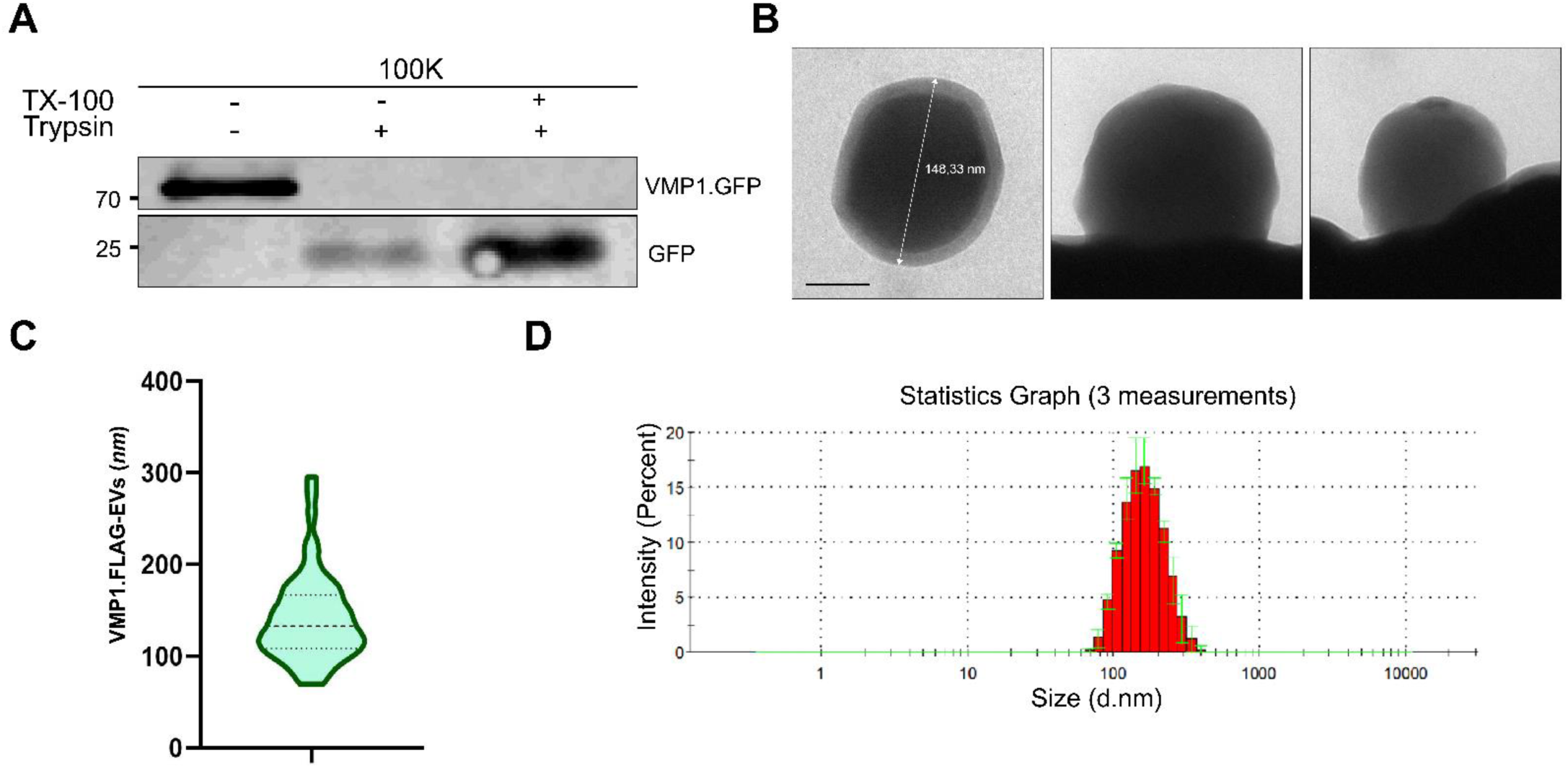
VMP1 is localized to the membrane of VMP1-EVs. (A) Immunoblot analysis with anti-GFP of VMP1.GFP EVs either untreated or treated with 100 μg/mL trypsin, with or without 1% Triton X-100 (TX-100), for 30 minutes at 37 °C. TX-100 was used as a positive control for trypsin accessibility. Protease sensitivity was assessed to determine membrane association of VMP1. Representative data from n = 3 biologically independent experiments are shown. (B) Representative TEM image of VMP1.FLAG–positive EVs isolated from the CM of HEK293T cells and purified by immunoprecipitation using anti-FLAG magnetic beads. Vesicles range in size from 150 to 200 nm. Scale bar: 50 nm. (C) Violin plot showing the size distribution of VMP1.FLAG-EVs based on TEM measurements (n = 25 vesicles). (D) DLS profile showing the intensity-based size distribution of VMP1.FLAG-EVs purified from CM (n = 3 biologically independent experiments).

In order to selectively isolate the VMP1-EVs, we used the fusion protein VMP1.FLAG, which has the tag binded to the VMP1 carboxyterminal aminoacid. Then, we immunoprecipitated VMP1.FLAG-EVs from HEK293T cell CM using anti-FLAG magnetic beads. TEM of the eluates revealed vesicular structures 110–200 nm in diameter (Figure 2B, C). DLS confirmed a homogeneous size distribution (Figure 2D). These data demonstrate that VMP1 is embedded in the EV membrane with its C-terminal domain facing outward.

### 2.3. VMP1 secretion depends on autophagy

Given VMP1’s essential role in autophagy, we examined whether its secretion is regulated by mTOR signaling. Nutrient starvation or PP242 treatment of VMP1.FLAG-expressing HEK293T cells reduced VMP1.FLAG secretion in EVs (Figure 3A). In contrast, chloroquine treatment increased VMP1.FLAG secretion, linking VMP1 release to autophagic flux. (Figure 3B).

**Figure 3.**
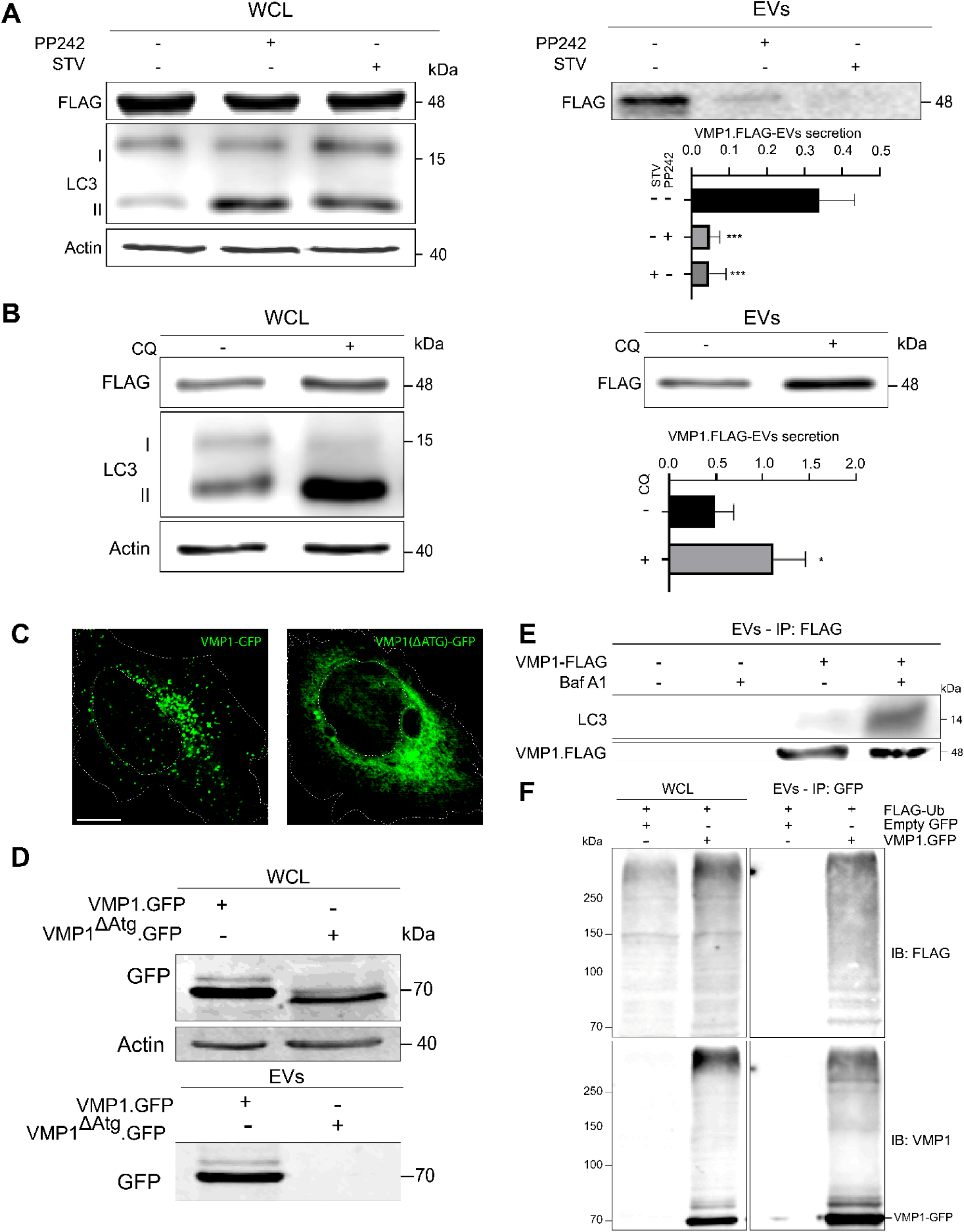
VMP1 Secretion is Dependent on Autophagy. (A) HEK293T cells transfected with VMP1.FLAG were treated for 16 h with either starvation medium (STV) or the mTOR inhibitor PP242 (1 µM). Immunoblot analysis of WCL was performed using antibodies against FLAG, LC3, and actin (loading control). EV fractions were analyzed by immunoblotting with anti-FLAG to detect secreted VMP1.FLAG. The bar chart shows the mean ± SEM of VMP1.FLAG levels in EVs, normalized to VMP1.FLAG levels in WCL relative to actin. Data represent mean ± SEM from n = 3 biologically independent experiments. Statistical analysis was performed using one-way ANOVA; *p < 0.01 versus control. (B) HEK293T cells transfected with VMP1.FLAG were treated with chloroquine (CQ, 50 µM) for 16 h to block autophagic flux. Immunoblot analysis of WCL was performed against FLAG, LC3, and actin (loading control). EV fractions were analyzed by immunoblotting with anti-FLAG to detect VMP1.FLAG. The bar chart shows the mean ± SEM of VMP1.FLAG levels in EVs, normalized to VMP1-FLAG levels in WCL relative to actin. Data represent mean ± SEM from n = 3 biologically independent experiments. Statistical significance was determined using one-way ANOVA; *p < 0.01 versus control. (C) Confocal images of HeLa cells transfected overnight with VMP1.GFP or VMP1ΔAtg.GFP. Direct GFP fluorescence was used to visualize protein localization. Scale bar: 10 µm. (D) HEK293T cells transfected with VMP1.GFP or VMP1ΔAtg.GFP. Immunoblot analysis of WCL with anti-GFP and antiactin (loading control) and of EV fractions with anti-GFP. (E) HEK293T cells transfected with VMP1.FLAG were treated overnight with BafA1 (10 nM; 16 h). EVs were immunoprecipitated using anti-FLAG magnetic beads, followed by immunoblot analysis for FLAG and LC3. (F) Immunoblot analysis of WCL and EV fractions from HEK293T cells co-transfected with FLAG-Ub and either empty-GFP or VMP1.GFP. Immunoprecipitation was performed using magnetic beads conjugated with anti-GFP, followed by immunoblotting for VMP1 (N-terminal) and FLAG. The VMP1 immunoblot shows multiple bands above 73 kDa in eluates from both total lysates and EVs, consistent with ubiquitinated VMP1.

Using the autophagy-deficient mutant VMP1ΔAtg.GFP, which is retained in the ER, we found no secretion into EV fractions (Figure 3C, D), indicating that autophagosome formation is required. To further link VMP1 secretion to autophagy, VMP1.FLAG–positive EVs were immunopurified from cells treated with BafA1. BafA1 increased LC3-II content within VMP1.FLAG EVs (Figure 3E), indicating enrichment of autophagic cargo under lysosomal inhibition. The corresponding immunoblots of WCL are shown in Figure A1.

Finally, given that VMP1 is ubiquitinated early in autophagosome formation and remains ubiquitinated throughout autophagy, EVs from cells co-expressing VMP1.GFP and FLAG.Ub were immunoisolated using anti-GFP beads. Ubiquitinated VMP1 was detected in EVs from VMP1.GFP and FLAG-Ub coexpressing cells (Figure 3F), consistent with secretion during autophagy.

### 2.4. Endogenous VMP1 secretion requires autophagosome formation and is induced by lysosomal inhibition

We tested whether lysosomal inhibition stimulates endogenous VMP1 secretion and whether autophagosome formation is required. In WT MEFs, BafA1 treatment increased VMP1 secretion into EV fractions, whereas Atg5 KO MEFs displayed markedly reduced VMP1-EV release (Figure 4A).

**Figure 4.**
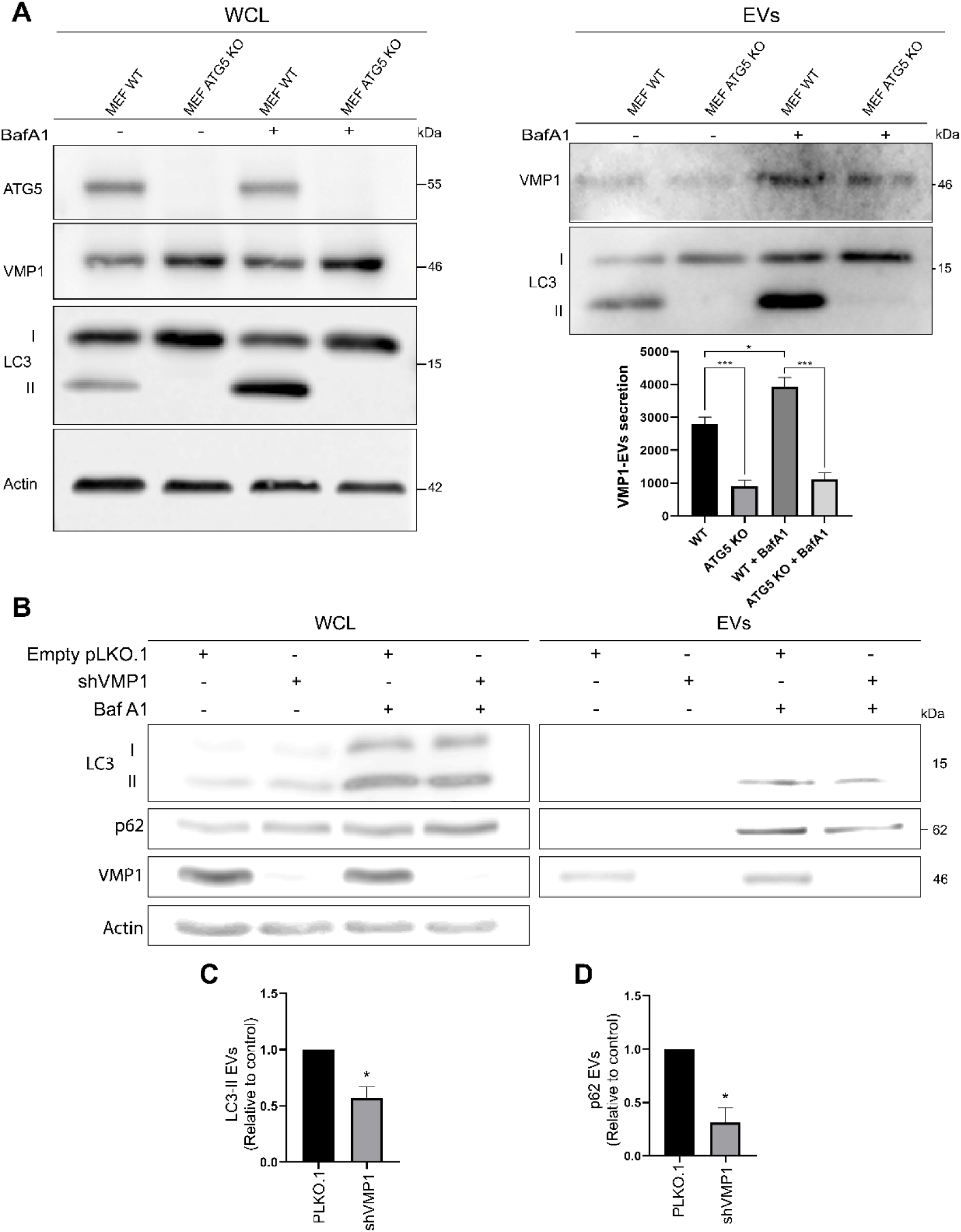
Endogenous VMP1 Secretion Requires Autophagosome Formation and Is Induced by Lysosomal inhibition. (A) VMP1 is detected in EVs independently of ATG5. EVs were isolated from wild-type (WT) and Atg5 knockout (Atg5 KO) mouse embryonic fibroblasts (MEFs) either untreated or treated with BafA1 (100 nM, 16 h). Immunoblot analysis of EV and WCL fractions was performed using antibodies against VMP1 (C-terminal) and LC3. ATG5 and actin were used as genotype and loading controls, respectively, in WCLs. The bar chart shows the mean ± SEM of VMP1 levels in EVs, normalized to VMP1 levels in WCL relative to actin. Data represent mean ± SEM (n = 3 independent experiments). Statistical analysis was performed using two-way ANOVA; *p < 0.05, **p < 0.01, ***p < 0.001. (B) MIA PaCa-2 cells with stable VMP1 KD (shVMP1) were treated overnight with BafA1 (20 nM, 16 h). WCL and EV fractions were analyzed by immunoblot for p62, VMP1 (N-terminal), LC3, and actin (loading control for WCL). (C) The bar chart shows the mean and SEM of LC3-II levels in EVs, normalized to LC3-II in WCL and actin, relative to control. p < 0.05 by Welch’s t-test vs. control. n= 3 independent experiments. (D) The bar chart shows the mean and SEM of p62 levels in EVs, normalized to p62 in WCL and actin, relative to control. p < 0.05 by Welch’s t-test vs. control. n= 3 independent experiments.

Given that lysosomal stress enhances p62 secretion via SA, we evaluated whether VMP1 contributes to this process using a MIA PaCa-2 cell line with constitutive VMP1 knockdown (VMP1 KD cells). In VMP1-expressing cells, BafA1 treatment induced the secretion of LC3-II and p62 in the EV fraction. In contrast, VMP1 KD cells showed a marked reduction in the secretion of both LC3-II and p62 (Figure 4B-D), indicating that VMP1 plays a critical role in the stress-induced release of autophagy-related cargos.

### 2.5. VMP1 secretion is upregulated in pancreatic acinar cells under lysosomal stress and experimental acute pancreatitis

Differentiated AR42J cells treated with BafA1 showed increased VMP1 secretion into EV fractions, accompanied by elevated LC3 and p62 levels (Figure 5A), indicating activation of secretory autophagy (SA) under lysosomal stress in pancreatic cells. The high LC3-II levels in whole-cell lysates (WCL) also reflect the blockade of autophagic flux caused by lysosomal inhibition.

**Figure 5.**
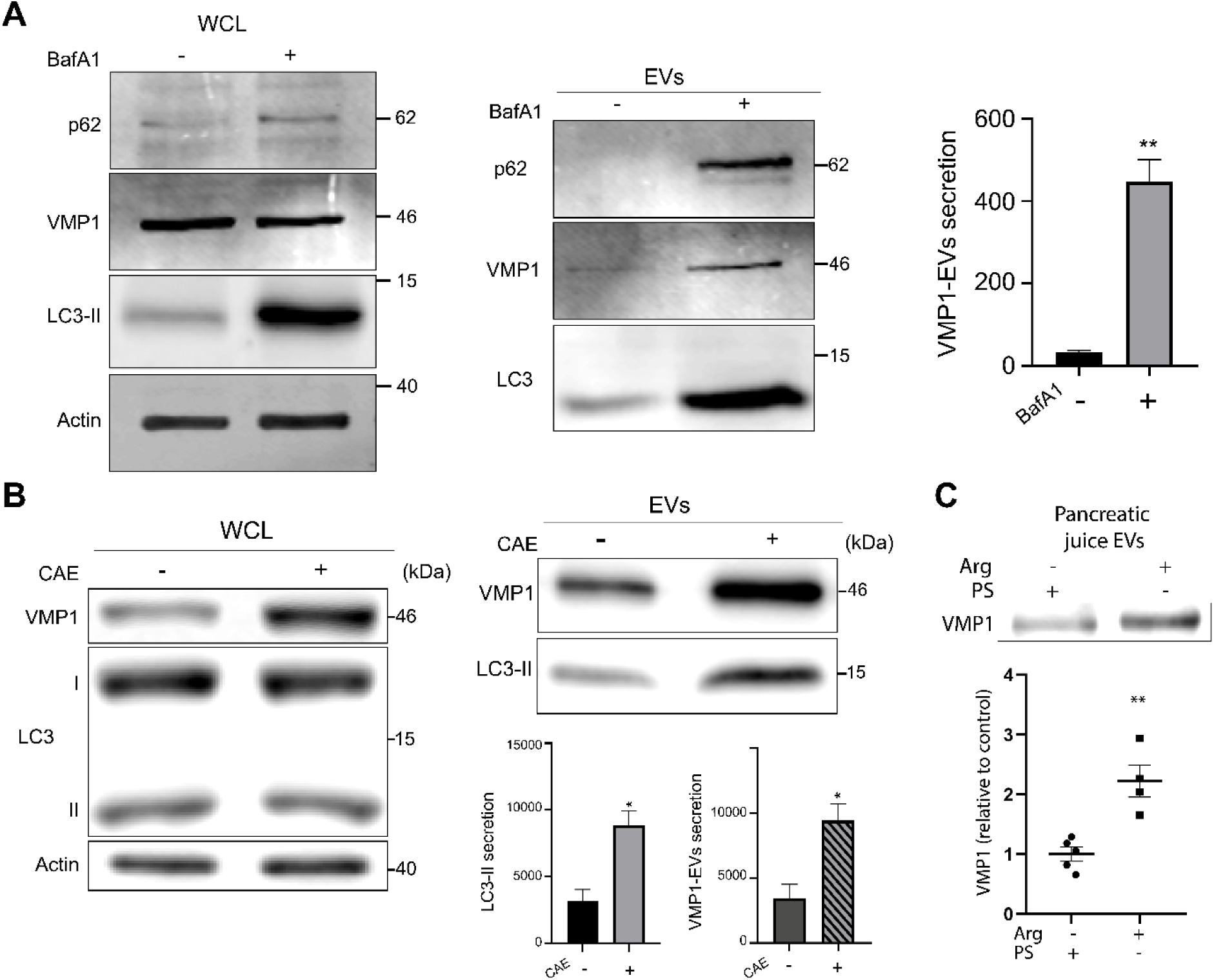
VMP1 Secretion Is Upregulated in Pancreatic Acinar Cells under Lysosomal Stress and Experimental Acute Pancreatitis. (A) AR42J cells were treated with BafA1 (20 nM, 16 h) or left untreated. Immunoblot analysis of WCLs and extracellular vesicle (EV) fractions was performed to detect VMP1 (C-terminal), p62, and LC3. Actin was used as a loading control for WCLs. The bar chart shows the mean ± SEM of VMP1 levels in EVs, normalized to VMP1 levels in WCL relative to actin. Bars represent the mean ± SEM from three independent experiments. Statistical analysis was performed using Welch’s t-test; **p < 0.01. (B) Cellular model of AP: AR42J cells were treated with caerulein (CAE, 7.4 μM, 16 h) or left untreated. Immunoblot analysis of WCLs and EV fractions was performed using anti-VMP1 (C-terminal) and anti-LC3 to assess VMP1 and LC3 levels in EV fractions. Actin was used as a loading control for WCLs. Bar charts show mean ± SEM of LC3-II and VMP1 in EVs, normalized to WCL levels of the same protein relative to actin. p < 0.05 by Welch’s t-test vs. control. n= 3 independent experiments. Bars represent the mean ± SEM from three independent experiments. Statistical analysis was performed using Welch’s t-test; *p < 0.05. (C) Animal model of AP: Immunoblot of VMP1 using anti-VMP1 (C-terminal) in the EV fraction obtained from PJ of rats treated with Arg or physiological solution (PS). The graph shows the quantification of VMP1 levels relative to the control. **: p < 0.01 by unpaired t-test vs. control. n = 5 for PS group, n = 4 for Arg group.

In a cellular model of acute pancreatitis (AP) induced by caerulein (CAE, 7.4 μM), EV fractions from treated AR42J cells contained higher levels of VMP1 and LC3-II than untreated controls, demonstrating that SA can be triggered by pancreatitis. The increased VMP1 levels in WCL indicate that experimental AP induces VMP1 expression (Figure 5B).

In vivo, EV fractions purified from pure pancreatic juice (PJ) of Arg-induced AP rats showed significantly higher VMP1 secretion compared to controls (Figure 5C). These findings identify a SA pathway in which upregulated VMP1 during pancreatitis is released via EVs.

## 3. Discussion

Autophagy is a tightly regulated catabolic process that enables the selective degradation of dysfunctional or superfluous cellular components, thereby maintaining cellular homeostasis. While autophagy is classically recognized for its role in cytoplasmic quality control, recent studies have elucidated other non-canonical functions, including participation in secretory pathways collectively termed SA [2]. These pathways encompass both conventional and unconventional protein secretion, the release of EVs, and secretory lysosome exocytosis [23,24].

Here, we demonstrate that VMP1, an autophagy-related transmembrane protein, is secreted within the EV fraction as an integral membrane component. VMP1 secretion is modulated by the autophagic status, being increased by lysosomal blockade and reduced by mTOR inhibition. It requires the expression of the core autophagy protein ATG5, and VMP1-EVs also carry LC3-II, indicating their co-secretion. We also show that VMP1 expression critically influences the secretion of the autophagy-related proteins LC3 and p62. These findings identify VMP1 as a novel contributor to SA. Moreover, we show that different stressors induce VMP1 secretion from pancreatic acinar cells, and that acute pancreatitis in rats promotes the release of VMP1-EVs into pancreatic juice, underscoring a key link between VMP1 secretion and the cellular response to acute pancreatitis.

VMP1 was detected in the EV fraction regardless of the tag or promoter used for its expression (Figure 1), including when expressed under a strong promoter such as CMV in pEGFP and pcDNA4-B plasmids, or under a moderate promoter such as PGK in the pLenti plasmid. Moreover, endogenous VMP1 was also detected in the EV fraction, and its secretion increased under cellular stress conditions (Figures 4 and 5). The secretion of VMP1 as part of the EV fraction represents a completely novel finding. This observation is consistent with other reports describing the secretion of autophagy-related molecules via EVs, such as LC3, ATG16L1, and LAMP2 (Pallet et al., 2013; Sirois et al., 2012), as well as p62, NBR1, WIPI2, and OPTN (Hessvik et al., 2016).

Regarding the size of these VMP1-EVs, using confocal microscopy we visualized GFP-tagged VMP1-EVs (VMP1.GFP-EVs) in conditioned medium, with sizes consistent with EVs (<1 μm; Figure 1). TEM and DLS further confirmed a homogeneous EV population, ranging from 110 to 200 nm (Figure 2). These data are consistent with those EV diameters described in MISEV2023 guidelines [4].

VMP1 is a transmembrane protein [Dusetti et al., 2002] localized to the autophagosomal membrane [Grasso et al., 2011; Renna et al., 2023]. As expected from its topology during autophagosome biogenesis, our results show that VMP1 is embedded in the EV membrane. Protease protection assays and TEM analyses of immunopurified VMP1-EVs confirmed that the protein’s hydrophilic C-terminus peptide is exposed on the EV surface (Figure 2), suggesting a novel role of VMP1 as an EV-integrated membrane protein.

The secretion of overexpressed tagged VMP1 was reduced upon inhibition of mTOR by both the pharmacological agent PP242 and starvation (Figure 3). Our results are consistent with other reports showing decreased EV output under such conditions [25,26]. On the other hand, inhibition of lysosomal degradation using chloroquine enhanced the secretion of overexpressed VMP1. Similar results were reported by other authors indicating that chloroquine treatment increases EV secretion [27] and autophagy-related proteins [28]. In addition, we found that the secretion of endogenous VMP1 in EVs also increases upon inhibition of lysosomal degradation, in this case with BafA1 (Figure 4), which is in agreement with previous reports describing BafA1 as an enhancer of EV secretion [29,30]. The secretion of VMP1, an autophagy-related protein, was found to depend on ATG5, a core component of the autophagy machinery. VMP1 secretion was significantly reduced in Atg5-deficient cells, in line with other studies showing that EV secretion depends on ATG5 expression [8,31,32]. Moreover, by immunoprecipitation, we detected LC3-II in VMP1-EVs, further supporting the notion that components of the autophagic machinery are directly associated with these vesicles and may contribute to their biogenesis and release. As a whole, these results indicate a clear relationship between VMP1 secretion and the autophagy machinery, suggesting that VMP1-EVs are released by a SA pathway.

Supporting the hypothesis of VMP1 being part of a novel SA pathway, we build on prior work showing that the C-terminal domain of VMP1 is required for autophagic flux [11,12]. In line with this, VMP1ΔAtg.GFP was not detected in EVs (Figure 3), indicating that the C-terminus is likewise necessary for entry into the secretory route. Moreover, we recently showed that VMP1 is ubiquitinated during autophagy [14]. Notably, here we find that VMP1 is also ubiquitinated when associated with EVs, reinforcing a mechanistic connection—and potential crosstalk—between autophagy and VMP1–EV secretion. Notably, we found that VMP1 is not only incorporated into EVs, but its expression also positively influences the secretion of two autophagy-related proteins, LC3 and p62 (Figure 4). Under lysosomal blockade conditions induced by BafA1, the secretion of LC3 and p62 was significantly reduced in VMP1 KD cells, indicating that VMP1 contributes to the release of these autophagic proteins. A schematic model illustrating the proposed pathway of SA involved in VMP1-EV secretion according to the main findings of this manuscript is shown in Figure 6.

**Figure 6.**
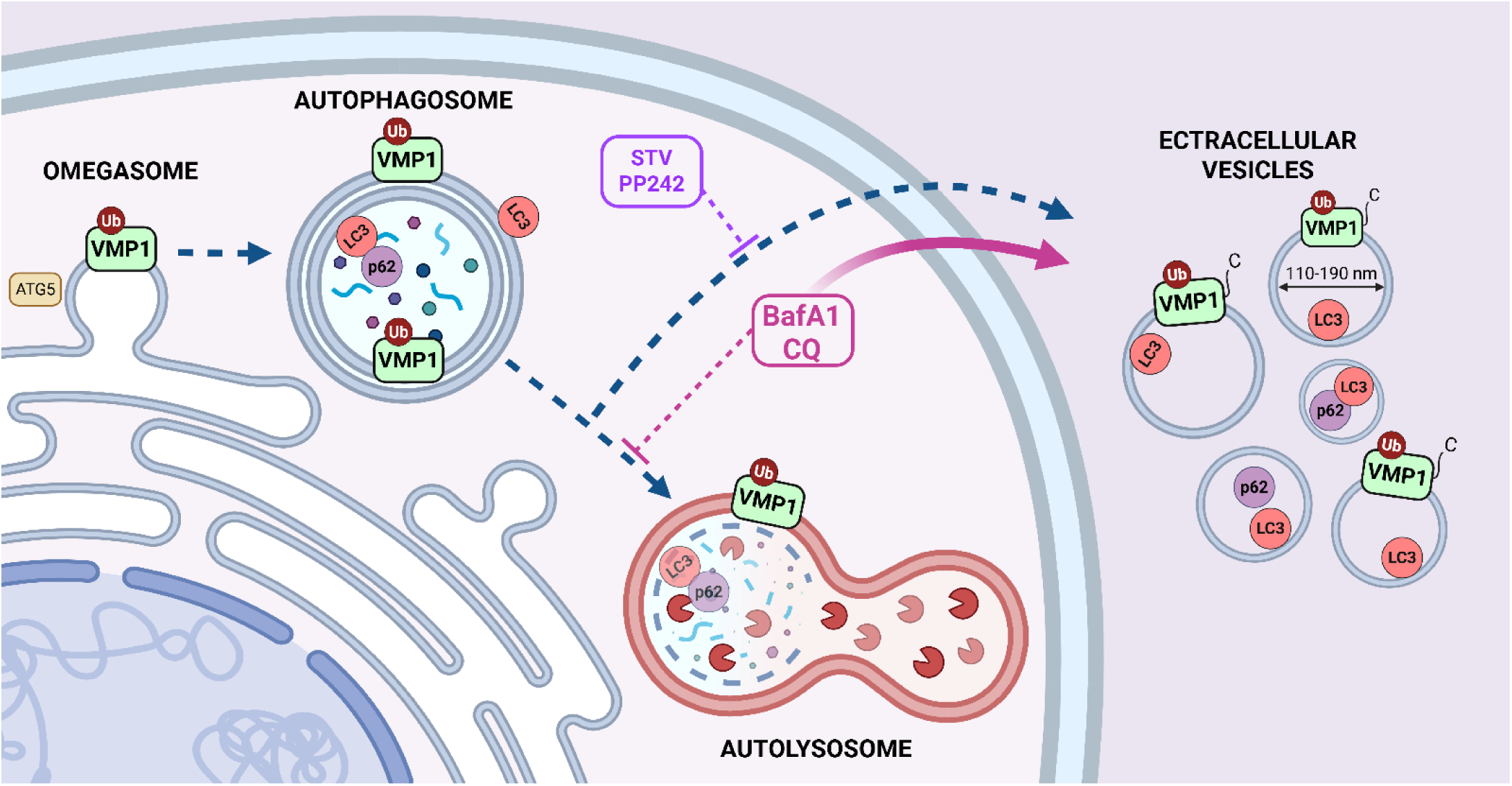
Proposed model of VMP1-EV release in SA responses to stress. Schematic model. VMP1 is secreted as a membrane component of extracellular EVs, ubiquitinated, with its C-terminal domain exposed on the outer surface. VMP1-EVs also contain LC3-II and are mainly 110–190 nm in diameter. VMP1-EV secretion depends on ATG5, is decreased by mTOR inhibition (via starvation or PP242 treatment), and is increased under lysosomal blockade (via chloroquine or BafA1). VMP1 is involved in LC3 and p62 secretion. Figure created with Biorender.com.

Importantly, SA is activated by a variety of cellular stressors, including endoplasmic reticulum (ER) stress, accumulation of misfolded proteins, and impaired intracellular trafficking— conditions closely linked to the pathogenesis of AP (Jahangiri et al., 2022; Zahoor and Farhan, 2018; Gonzalez et al., 2020). In AP, premature activation of digestive zymogens within pancreatic acinar cells initiates a cascade of tissue injury and inflammation. We previously demonstrated that VMP1 is markedly upregulated in pancreatic acinar cells during AP, where it promotes zymophagy—a selective form of autophagy that mitigates zymogen-induced cytotoxicity—and mitophagy, thereby pre-serving cellular energy homeostasis and preventing disease progression (Grasso et al., 2011; Vanasco et al., 2021; Boggio et al., 2025).

In the present study, we demonstrate for the first time that AP-mimicking conditions in pancreatic acinar cells not only induce a significant increase in VMP1 expression but, importantly, also stimulate its secretion via EVs (Figure 5). Moreover, we investigated VMP1 secretion in vivo using a widely accepted animal model of acute pancreatitis (Hegyi et al., 2004), in which we previously reported VMP1 upregulation in pancreatic acinar cells (Vaccaro et al., 2003). Notably, VMP1 was detected in the EV fraction isolated from PJ. Following Arg-induced AP in rats, VMP1 secretion was significantly increased in PJ, confirming its induction under pathological conditions.

These findings reveal a novel secretory role for VMP1 in response to pancreatitis-associated stress and highlight its potential relevance in pancreatic disease (Figure 7).

**Figure 7.**
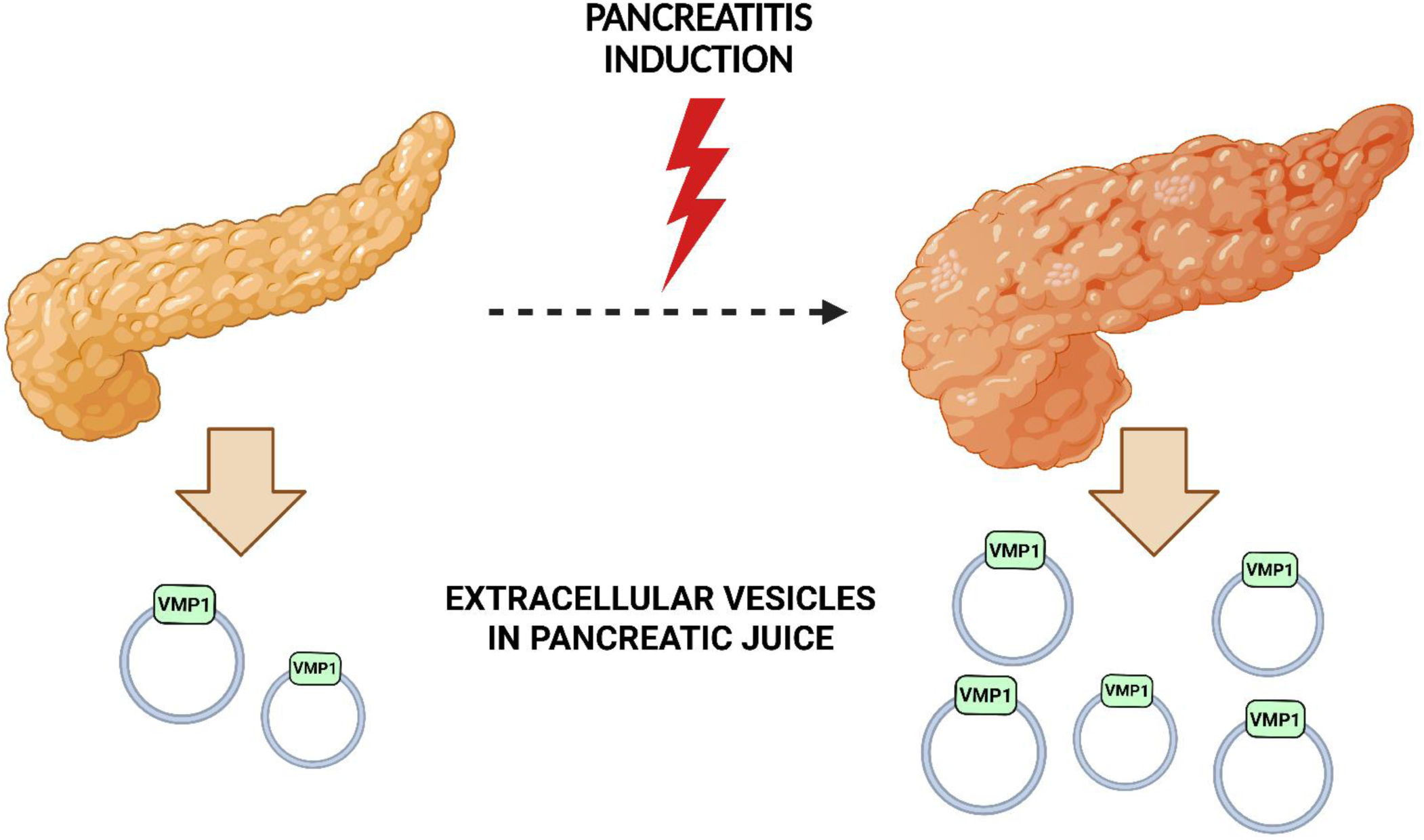
Cartoon showing VMP1-EV release during pancreatitis. VMP1 is secreted in EVs into PJ, with levels markedly increased upon pancreatitis induction. Figure created with Biorender.com.

## 4. Materials and Methods

### 4.1 Animal Model of AP

AP was induced in male and female Wistar rats (8–10 weeks old) with Arginine (Arg) by intraperitoneal injection of L-arginine hydrochloride, a well-established model of pancreatic injury mediated by metabolic and oxidative stress. Animals received two i.p. injections of 250 mg Arg per 100 g body weight, prepared as a 20% solution in sterile saline (pH adjusted to 7.0), with a 1 h interval between doses. Rats had ad libitum access to food and water between injections. Animals were euthanized at indicated time points for histological and biochemical analysis of pancreatic tissue. Control animals received equivalent volumes of sterile saline following the same injection schedule. All animal procedures were approved by the Animal Care and Research Committee of the School of Pharmacy and Biochemistry, University of Buenos Aires (REDEC 2024 1664 E UBA DCT FFYB), and were conducted in accordance with the International Guiding Principles for Biomedical Research Involving Animals.

### 4.2 Cell Culture, Transfections, and Differentiation

HEK293T (RRID:CVCL_0063), HeLa (RRID:CVCL_0030), MIA PaCa-2 (RRID:CVCL_0428) and AR42J (RRID:CVCL_0143) cell lines were obtained from the American Type Culture Collection. Wild-type (WT) (RRID:CVCL_C1M8) and Atg5-knockout MEFs (RRID:CVCL_0J75) were generously provided by Dr. María I. Colombo (Universidad Nacional de Cuyo, CONICET, Argentina). MIAPaCa-2 pancreatic cancer cells with stable knockdown of VMP1, achieved via shRNA expression, and vector control cells were used to evaluate VMP1 expression. All cell lines were cultured in Dulbecco’s Modified Eagle Medium (Biological Industries) supplemented with either 10% or 20% fetal bovine serum (Natocor), 200 mM L-alanyl-L-glutamine dipeptide (Gibco™), 100 U/mL penicillin, and 100 μg/mL streptomycin (Biological Industries). Cells were maintained at 37°C in a humidified 5% CO₂ incubator. Mycoplasma contamination was routinely assessed by DAPI staining (monthly) and by PCR (biannually). HEK293T cells were transfected at 60–70% confluence using polyethyleneimine (Polysciences, cat. no. 24765-1). Culture medium was replaced with fresh conditioned medium (CM) 5 h post-transfection. AR42J cells were differentiated by incubation with 100 nM dexamethasone for 48 h prior to experimental procedures.

### 4.3 Treatments

For amino acid deprivation, cells were washed twice with fasting medium (140 mM NaCl, 1 mM CaCl₂, 1 mM MgCl₂, 5 mM glucose, 1% BSA, and 20 mM HEPES, pH 7.4) and incubated in the same medium for 16 h to collect CM. PP242 (Santa Cruz Biotechnology) was dissolved in DMSO to prepare a 1 mM stock solution, and cells were treated with 1 µM PP242 for 16 h. Chloroquine diphosphate (Sigma-Aldrich) was dissolved in phosphate-buffered saline (PBS) to obtain a 10 mM stock solution; cells were treated with 50 µM chloroquine for 16 h. Bafilomycin A1 (Santa Cruz Biotechnology; cat. no. sc-201550A) was prepared as a 10 µM stock solution in DMSO and applied at final concentrations, according to cell types, of 10 nM, 20 nM or 100 nM for 16 h. For hyperstimulation of cholecystokinin receptors, differentiated AR42J cells were treated with 7.4 µM caerulein (Sigma-Aldrich) for 16 h.

### 4.4 Expression Vectors

The plasmids pEGFP-N1-hVMP1 (VMP1.GFP), pcDNA4-B–hVMP1–V5/His (VMP1.V5), and pLenti–hVMP1–FLAG (VMP1.FLAG) were generated as previously described in Renna et al., 2023 [14]. The pLKO.1-Puromycin vector containing a pre-designed shRNA sequence targeting human VMP1 (5′-GCAATGAACAAGGAACATCAT-3′; catalog number RHS3979-98821402) was used to generate MIAPaCa-2 VMP1 knockdown (VMP1 KD) cells. A deletion construct lacking the C-terminal 18 amino acids of VMP1 (residues 388–406), termed VMP1ΔAtg, was cloned into the pENTR1A-GFP-N2 vector (Addgene plasmid #19364) using BamHI and EcoRI restriction sites, based on the human VMP1 reference sequence (NM_030938.5). This entry clone was recombined with the pLenti PGK puro destination vector (Addgene plasmid #19068) using Gateway™ LR Clonase™ II enzyme mix (Thermo Fisher Scientific; cat. no. 11791-020). All constructs generated in this study were verified by DNA sequencing. The FLAG–Ubiquitin (FLAG.Ub) plasmid was generously provided by Dr. Simona Polo (University of Milan). We used pEGFP-N1 (empty GFP) as a control.

### 4.5 Generation of Conditioned Culture Medium

To generate exosome-depleted CM, DMEM (Biological Industries) supplemented with 200 mM L-alanyl-L-glutamine dipeptide (Gibco™) and 30% Fetal Bovine Serum (Natocor) was ultracentrifuged at 120,000 × g for 16 h using a Beckman XL-90 ultra-centrifuge. The resulting supernatant was filtered through 0.22 µm pore-size filters and transferred to sterile containers. Penicillin (100 U/mL) and streptomycin (100 µg/mL) (Biological Industries) were added to the filtered medium.

### 4.6 Isolation of EVs from Conditioned Culture Medium

CM was collected and subjected to a series of differential centrifugation steps to re-move cells and debris: 300 × g for 15 min, followed by 2,000 × g for 20 min at 4°C. The supernatant was then centrifuged at 16,500 × g for 20 min at 4°C. Subsequently, the supernatant was ultracentrifuged at 120,000 × g for 2 h at 4 °C using a Beckman XL-90 ultracentrifuge. The resulting pellet was washed with cold PBS and ultracentrifuged again at 120,000 × g for 1.5 h at 4°C. The final EV pellet was resuspended in PBS and used for downstream applications, including immunoblotting, immunofluorescence, or immunoprecipitation.

### 4.7 Isolation of EVs from Rat Pancreatic Juice

Pancreatic juice (PJ) collected from experimental animals was centrifuged at 300 × g for 10 min at 4°C, followed by 2,000 × g for 10 min at 4°C to remove cellular debris. The supernatant was then centrifuged at 10,000 × g for 30 min at 4°C. This fraction was diluted in PBS and subjected to ultracentrifugation at 100,000 × g for 2 h at 4°C using a TFT 80.2 rotor in a Sorvall WX+ Ultra Series ultracentrifuge (Thermo Scientific). The pellet was washed with PBS and subjected to a second ultracentrifugation step at 100,000 × g for 1.5 h at 4°C. The final EV pellet was resuspended in PBS for subsequent immunoblotting analysis.

### 4.8 Immunoblot Analysis

Following treatments and transfections, cells were lysed in buffer containing 50 mM Tris-HCl (pH 7.4), 250 mM NaCl, 25 mM NaF, 2 mM EDTA, 0.1% Triton X-100, and protease inhibitors (Thermo Scientific™ Pierce). EV pellets were lysed in the Laemmli buffer. For LC3 detection, cells were lysed in 50 mM Tris-HCl (pH 8.0), 150 mM NaCl, 1% Triton X-100, and protease inhibitors. Protein concentration was quantified using the BCA assay (Pierce), and equal amounts of protein were resolved by SDS-PAGE and transferred to polyvinylidene fluoride (PVDF) membranes (0.22 μm pore size; Millipore). Membranes were blocked in Odyssey Blocking Buffer (LI-COR) for 1 h at room temperature and incubated overnight at 4°C with the appropriate primary antibodies. Primary antibodies included: anti-VMP1 (N-terminal) (1:1,000; rabbit mAb #12978, RRID:AB_2798077, Cell Signaling Technology), anti-VMP1 (C-terminal) (1:1,000; rabbit mAb #12929, RRID:AB_2714018, Cell Signaling Technology), anti-actin (1:2,000; rabbit polyclonal A2066, RRID:AB_476693, Sigma– Aldrich), anti-DYKDDDDK tag (1:6,000; mouse mAb #8146, RRID:AB_10950495, Cell Signaling Technology), anti-α-tubulin (1:4,000; mouse mAb T5168, RRID:AB_477579, Sigma–Aldrich), anti-SQSTM1/p62 (1:1,000; rabbit mAb #8025, RRID:AB_10859911, Cell Signaling Technology), anti-CD63 (1:1,000; mouse mAb ab231975, Abcam), anti-GFP (1:500; mouse monoclonal antibody 3E6, RRID:AB_221568, Invitrogen), anti-ATG5 (1:1000; rabbit monoclonal, #12994, RRID:AB_2630393, Cell Signaling Technology) and anti-V5 (1:5000; mouse monoclonal, R960-25, RRID:AB_2556564, Invitrogen). After primary antibody incubation, membranes were washed four times with PBS containing 0.1% Tween-20 (PBS-T), then twice with PBS. Membranes were then incubated for 2 h at room temperature with secondary antibodies: IRDye® 680LT goat anti-rabbit IgG (1:15,000) or IRDye® 800CW goat anti-mouse IgG (1:10,000; LI-COR), diluted in Odyssey Blocking Buffer. Following incubation, membranes were washed again and visualized using the Odyssey® SA system (LI-COR). For LC3 detection, membranes were blocked for 1 h at room temperature in 1% bovine serum albumin (BSA) in TBS containing 0.1% Tween-20 (TBS-T), then incubated with anti-LC3B (1:500; rabbit mAb #3868, RRID:AB_2137707, Cell Signaling Technology) for 24 h at 4°C. Membranes were washed, then incubated with HRP-conjugated anti-rabbit IgG (1:2,500; Amer-sham NA934, GE Healthcare) in 1% BSA in TBS-T. Detection was performed using Pierce ECL Plus (Cat# 32134, Thermo Scientific) and visualized on a cDigit Blot Scanner (LI-COR). Band intensities were quantified using FIJI software. Densitometry values were normalized to actin and expressed as the mean ± SEM from three independent experiments.

### 4.9 Immunoprecipitation Assays

For anti-FLAG immunoprecipitation, EV pellets obtained by ultracentrifugation of conditioned culture media were resuspended in 300 μL of 1× PBS and transferred to 2 mL Eppendorf tubes. Samples were incubated with anti-FLAG® M2 magnetic beads (Sigma-Aldrich) for 2 h at 4°C with gentle agitation. Beads were then washed five times with PBS containing 0.02% Tween-20. Proteins were eluted by adding Laemmli buffer followed by heating at 95°C for 5 min. For anti-GFP immunoprecipitation, Pierce™ Protein G magnetic beads (Invitrogen) previously attached to the Green Flu-orescent Protein (GFP) tag antibody (RRID:AB_221568) were used. EV pellets were resuspended in 300 μL of 1× PBS and incubated with the beads for 1 h at 4°C with agitation. Beads were washed five times with PBS containing 0.02% Tween-20. VMP1.GFP-EVs were eluted by adding Laemmli buffer and heating at 95°C for 5 min.

### 4.10 Fluorescence Intensity of VMP1.GFP-EVs

HEK293T cells transfected with the VMP1.GFP plasmid were cultured, and conditioned media were collected and subjected to ultracentrifugation to isolate EVs. The resulting pellet was resuspended in PBS, and GFP fluorescence was quantified using a SpectraMax i3x microplate reader (Molecular Devices) at 510 nm. The normalized fluorescence intensity of VMP1.GFP-EVs was calculated using the formula: FI_EVs / FI_WCL, where FI_EVs represents the fluorescence intensity of EVs and FI_WCL represents the intensity of the corresponding whole cell lysate (WCL) from VMP1.GFP-transfected cells.

### 4.11 Dynamic Light Scattering (DLS)

DLS measurements were performed on EV pellets obtained from conditioned media of HEK293T cells transfected with the pLenti-Puro-PGK-hVMP1 plasmid. Measurements were carried out at 25°C using a Zetasizer Nano-ZSP system (Malvern Instruments, UK) equipped with a 633 nm He-Ne laser and a ZEN5600 digital correlator.

### 4.12 Transmission Electron Microscopy (TEM)

Following immunoprecipitation of VMP1.FLAG-EVs, samples were resuspended in PBS and fixed with 2% glutaraldehyde in 0.1 M phosphate buffer at 4°C. A small volume of the fixed sample was placed on a 200-mesh copper grid coated with LR white resin membrane (Sigma-Aldrich, L9774), washed eight times with ultrapure water (2 min each), and stained with 2% aqueous uranyl acetate (pH 7) for 3 min. Excess stain was removed with blotting paper, and grids were airdried for 15 min at room temperature. Imaging was performed on a Zeiss 109T transmission electron microscope operated at 80 kV, and images were acquired using a Gatan ES1000w digital camera.

### 4.13 Immunofluorescence

Cells were cultured on 15 mm round glass coverslips placed in 12-well plates. A total of 60,000 HeLa or 80,000 AR42J cells were seeded per well. The following day, cells were transfected with the indicated plasmids, if required, and subsequently incubated with control EVs or VMP1 containing EVs for 24 h. After treatment, cells were fixed with 3.6% paraformaldehyde in PBS for 20 min followed by permeabilization with 0.1% Triton X-100 in PBS for 5 min. Samples were then washed three times with PBS and blocked with 1% BSA in PBS for 1 h at room temperature. Coverslips were incubated overnight at 4°C in a humidified chamber with primary antibodies diluted in blocking solution. The following antibodies were used: anti-FLAG (1:500; mouse mAb, DYKDDDDK Tag (9A3), RRID:AB_10950495, Cell Signaling Technology) and anti-tubulin (1:1000; mouse mAb T5168, RRID:AB_477579, Sigma-Aldrich). After incubation, coverslips were washed three times with PBS and incubated with Alexa Flu-or-conjugated secondary antibodies (Alexa Fluor 488, 594, or 647; 1:800, Molecular Probes) for 2 h at room temperature in PBS with 1% BSA. Coverslips were mounted with polyvinyl alcohol (PVA) containing DABCO mounting medium. Confocal images were acquired using a Zeiss LSM 880 confocal microscope equipped with 20× (NA 0.8) and 63× (NA 1.4) Plan-Apochromat objectives and Zeiss ZEN Black software.

### 4.14 In Vivo Fluorescence Imaging of VMP1.GFP-EVs

HEK293T cells transfected with VMP1.GFP were used to generate EVs by ultracentrifugation of conditioned media. The EV-containing pellet was resuspended in PBS, and a drop of the suspension was placed on a microscope slide, covered with a glass coverslip, and sealed with nail polish. Samples were immediately imaged using a Zeiss LSM 880 confocal microscope with 20× or 63× Plan-Apochromat objectives.

### 4.15 Protease Protection Assay

VMP1.FLAG-EVs were isolated by differential centrifugation and resuspended in 60 μL PBS. The suspension was divided into three aliquots and incubated with: (i) PBS alone, (ii) 100 μg/mL trypsin in PBS, or (iii) 1% Triton X-100 with 100 μg/mL trypsin in PBS. Reactions were carried out for 30 min at 37°C with gentle mixing. Protease activity was halted by adding protease inhibitor cocktail and 2× Laemmli sample buffer. Samples were then subjected to immunoblotting.

### 4.16 Statistical Analysis

All quantitative data are presented as mean ± SEM. Representative images are shown from at least three independent experiments. Differences between groups of quantitative data were assessed by ANOVA, Student-Newman-Keuls post hoc test. P values below 0.05 (two tailed) were considered as statistically significant. Statistical analyses were performed using GraphPad Prism 8, as described in the corresponding figure legends.

## 5. Conclusions

We provide the first evidence that the autophagy-related transmembrane protein VMP1 is secreted via EVs through a SA pathway. VMP1-EV secretion is induced under conditions of lysosomal blockade and pancreatitis-associated stress and is dependent on autophagy. Our findings suggest that VMP1 not only mediates the degradation of damaged components through selective macroautophagy but also participates in non-canonical autophagic processes, including SA via EVs. This dual role may contribute to cellular adaptation and survival during stress and may be particularly relevant in the context of inflammatory diseases such as acute pancreatitis.

## Author Contributions

**Conceptualization**, MIV, MST and FJR; methodology, MST, MIV, FJR, MHL, FLM and CDG; validation, MIV and FJR; formal analysis, MST MIV, FJR, AR; investigation, MST, FJR, MHL, FLM, AR, TO and DAC; funding acquisition and resources AR, CDG and MIV; data curation, FJR and AR; writing—original draft preparation, MST, FJR and MIV.; visualization, MST, FJR and MIV.; supervision, MIV, FJR, AR and CDG; project administration, MIV. All authors have read and agreed to the published version of the manuscript.

## Institutional Review Board Statement

All animal procedures were approved by the Animal Care and Research Committee of the School of Pharmacy and Biochemistry, University of Buenos Aires (REDEC 2024 1664 E UBA DCT FFYB), and were conducted in accordance with the International Guiding Principles for Biomedical Research Involving Animals.

## Data Availability Statement

The raw data are available from the corresponding author upon reasonable request.

## Acknowledgments

We gratefully acknowledge Julieta Agustina Repetti, Carolina Verónica Vecino and Veronica Boggio for their valuable assistance with PJ collection in the rat model.

## Abbreviations

The following abbreviations are used in this manuscript:

AP: Acute Pancreatitis
Arg: Arginine
BafA1: Bafilomycin A1
CM: Conditioned Medium
DLS: Dynamic Light Scattering
EVs: Extracellular Vesicles
pEGFP-N1: Empty GFP
FLAG. Ub: FLAG–Ubiquitin
GFP: Green Fluorescent Protein
LC3: Microtubule-associated protein 1 light chain 3
P62: SQSTM1/p62 (sequestosome 1)
PJ: Pancreatic Juice
PBS: Phosphate-buffered saline
SA: Secretory autophagy
TEM: Transmission electron microscopy
VMP1: Vacuole Membrane Protein 1
VMP1ΔAtg: Construct lacking the C-terminal 18 amino acids of VMP1 (residues 388–406)
VMP1-EVs: VMP1-containing EVs
VMP1.FLAG-EVs: VMP1.FLAG-containing EVs
VMP1.FLAG: pLenti–hVMP1–FLAG
VMP1.GFP: pEGFP-N1-hVMP1
VMP1.GFP-EVs: VMP1.GFP-containing EVs
VMP1 KD: VMP1 knockdown
VMP1.V5: pcDNA4-B–hVMP1–V5/His
WCL: Whole cell lysate
WT: Wild-Type

## Appendix A

### Appendix A.1

**Figure A1.**
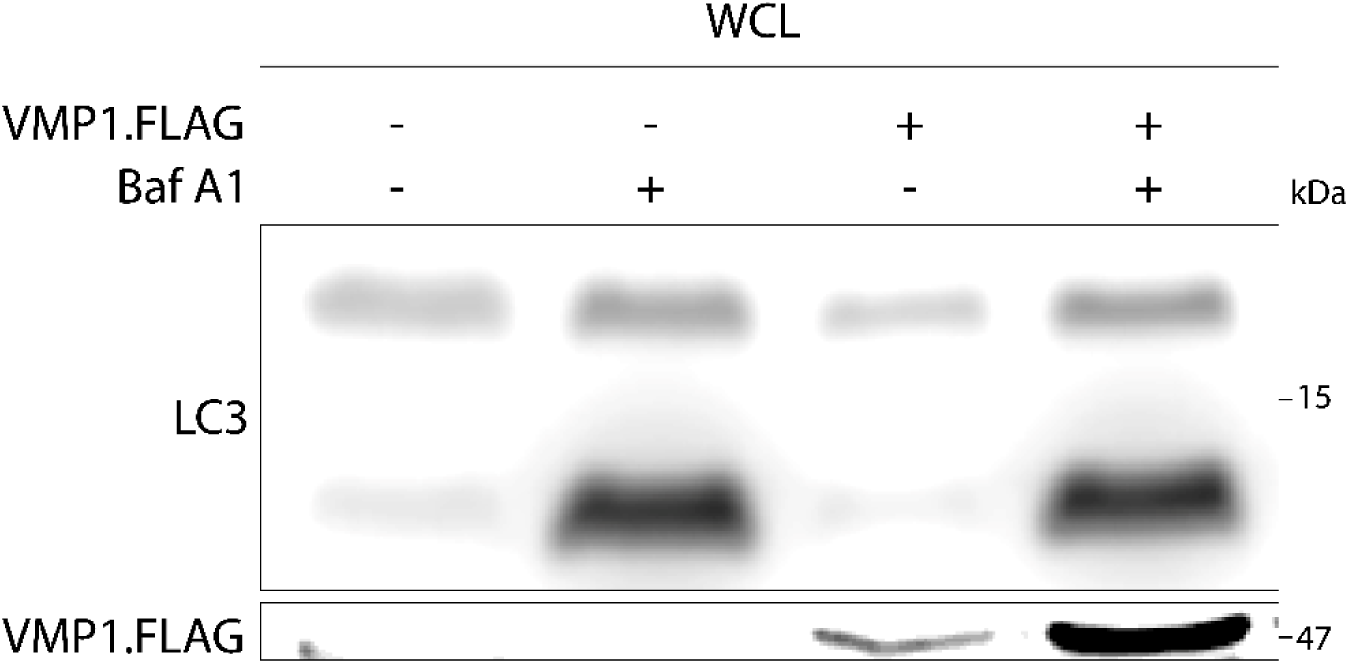
LC3 and VMP1-FLAG expression in WCL of HEK293T cells. Immunoblot with anti-LC3 and anti-FLAG of WCL corresponding to the IP shown in Figure 4E. HEK293T cells were transfected with VMP1.FLAG and treated with BafA1 (10 nM, 16 h).

## Disclaimer/Publisher’s Note

The statements, opinions and data contained in all publications are solely those of the individual author(s) and contributor(s) and not of MDPI and/or the editor(s). MDPI and/or the editor(s) disclaim responsibility for any injury to people or property resulting from any ideas, methods, instructions or products referred to in the content.

