## Supplementary figures and images for "Secretory Autophagy via VMP1-Containing Extracellular Vesicles in Pancreatic Stress Responses"

### Appendix A

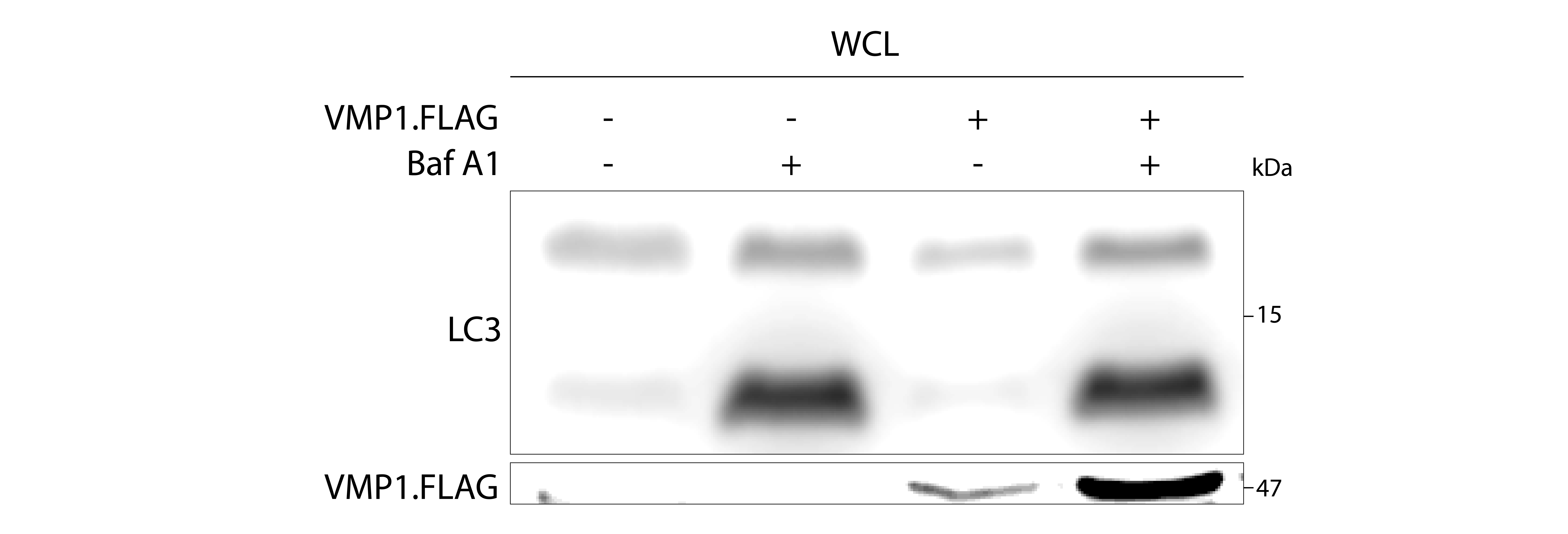
